# EpiTune: An Accurate Epitope Prediction Model with Mechanistic Insights

**DOI:** 10.64898/2026.08.12.744310

**Authors:** Chiril Calin, Dac-Trung Nguyen, B. Scott Perrin

## Abstract

The accurate prediction of b-cell epitopes facilitates vaccine development by identifying known antibodies for an antigen. Multiple epitope prediction models use protein language models to enable more accurate predictions with modest results. Here, we present EpiTune, a b-cell epitope prediction model that fine-tunes the underlying protein language model to deliver best-in-class predictions of linear epitopes and competitive predictions for confirmational epitopes. EpiTune achieves this performance from antigen sequence alone, and utilizes ESM-2’s RoPE architecture to fine-tune and infer on sequences longer than other sequence-based models currently available in the literature. EpiTune’s single-model architecture allows the model to determine the meaningfulness of sequence features for epitope prediction. This avoids the need for assigning importance to intermediates such as structure-based information, while still allowing a high degree of model interpretability.

## Introduction

B-cell lymphocytes are an integral part of the immune system, where they mediate mammalian immune response through antibody production^1^. To induce an immune response, the epitope (or antigenic determinant) of an antigen is recognized by the paratope on an antibody.

The experimental identification of epitopes through techniques such as co-crystallization, site-directed mutagenesis, hydrogen-deuterium exchange (HDX) is traditionally used for epitope mapping.^2^ Epitope prediction can help inform a direction for or confirm results from experimental epitope mapping, and as such, has been of great interest.

Epitopes are classified as either conformational or linear based on how they interact with the paratope. Conformational epitopes consist of discontinuous residues that are brought together by the antigen fold, whereas linear epitopes comprise continuous amino acid sequences. On average, epitopes span approximately 15 residues, typically including a continuous stretch of at least five residues that account for more than half of the epitope length.^3^ Although conformational epitopes are far more prevelant^4,5^ a large number of linear epitopes have been experimentally characterized due to their relative ease of identification. Consequently, predictive models for linear epitopes are generally more accurate, whereas the development of reliable conformational epitope prediction models would provide substantially greater practical value.

The epitope prediction problem can be modeled as a binary classification task, with a predictive model designating whether a given residue is an epitope or not. The performance of binary classification task models is generally measured by precision, recall, and the receiver-operator curve (ROC)^6^. A precision-recall (PR) curve is generated by assessing precision and recall scores at all possible thresholding classification points, and the integral of this curve provides the PR-AUC (area under the PR curve). The epitope prediction literature generally compares models by their ROC-AUC^7–9^, however, PR-AUC is more sensitive for imbalanced datasets where positive values are rare, such as in the case of epitope prediction.

Epitope prediction tools integrate machine learning principles and models into their mechanism. Among the earlier epitope prediction methods, models like BepiPred (2009) utilized Hidden Markov approaches^10^, which BepiPred 2.0^11^ (2017) built on. Most recently, several models have incorporated ESM-2 as part of their epitope prediction. ESM-2 is a protein language model trained on biological/protein sequence data^12^ and uses an encoder-only architecture alongside rotary embeddings like those used in RoFormer^13^. The transformer nature of the model allows sequence features to be represented robustly in high dimensional embedding space, and as such, many state-of-the-art models (BepiPred 3.0, DiscoTope, SEMA, ScanNet) use ESM-2’s embeddings. However, these approaches typically use ESM-2 as an embedding device only, and train separate machine learning models on its representations.

EpiTune takes advantage of ESM-2’s full transformer architecture by tuning directly on IEDB data^14^. By overlapping all epitopes onto their respective antigens, we fine-tune ESM-2 towards the epitope classification task, translating from unlabeled antigen sequences to sequences of binary epitope positional maps. In fine-tuning, we highlight that transfer learning of ESM-2’s general protein features is necessary for performant epitope prediction. We also compare model performance and reasoning at differing model sizes, tuning very large models such as ESM2-15B, and expand past the 1,024 sequence length cutoff point that other sequence ESM-based models limit themselves to^8,15^. Finally, we highlight that EpiTune’s single model architecture allows it to discover meaningful sequence features, and using sequences alone, demonstrate superior performance to models that require antigen structures for prediction.

## Methods

### Data Retrieval

Datasets were generated by pulling all human B-cell linear and conformational epitopes from IEDB as of Jul. 2025, totaling ∼180,000 linear epitopes across 8,208 antigen sequences, and ∼4,000 conformational epitopes across 379 antigen sequences. Epitopes were split between linear and conformational, and were paired with their respective antigens by UniProt ID, allowing for several epitopes to be mapped onto a single antigen. All assay records were compared for each epitope; epitopes with a greater amount of negative qualitative assays than positive were filtered out, resulting in ∼70,000 epitopes across 7,400 antigen sequences. This process was only performed for the linear dataset, as the conformational dataset only contained ∼400 negative assays out of 30,000 total assays. Extracted antigens varied from ∼30 to ∼33,000 AA in length, with a mean antigen sequence length of ∼850.

### Dataset Clustering Analysis

To assess the similarity between sequences, clustering using the cd-hit algorithm^4^ was performed at 80% similarity. Cd-hit found a total of ∼7,600 clusters, with 5% of clusters containing more than one sequence, and 0.4% of clusters containing more than 3 sequences. To ensure similar sequences were not present in training/test sets and to prevent overfitting, clusters were formed at 80% similarity and only one sequence was chosen per cluster, resulting in 7,714 total linear sequences and 345 conformational sequences. Then, to further reduce the linear dataset, MMseqs2^16^ clustering at 50% similarity was performed, and the representative sequence from each cluster was chosen, resulting in a final count of 6,284 linear sequences. The conformational dataset was not reduced further.

### Dataset Generation

Finally, both linear and conformational datasets were split according to a 90-10 train-test split, resulting in 566 linear and 35 conformational test set sequences. Validation datasets were not created as hyperparameter tuning was not used in the work. To assess performance of the structure-based epitope prediction models, structures were retrieved for each item in the linear and conformational test sets. Structures were extracted programmatically for each test set sequence:

1. Query UniProt API with IEDB parent antigen URL to get list of 3d structures available.
2. Identify whether an entry has RCSB PDB structures that contain, at minimum, all epitope positions identified on the antigen. If multiple structures contain all epitope positions identified on the antigen, return the structure with the highest resolution.
3. If entry does not contain PDB structures, or if all PDB structures available do not model the epitope positions, return AlphaFoldDB structure if available.

### Fine-Tuning Architecture

ESM-2 was used as the foundational model for fine-tuning EpiTune. We designed our training for epitope prediction as a machine classification task, with lowercase sequences being used as inputs, and 0/1 epitope status per residue position as our reference labels.

Sequences greater than 1,724 residues in length were removed from the training dataset due to the disproportionally increased computational requirements from longer sequences in a transformer architecture. This cutoff point represented a removal of only the top 10% of longest sequence lengths. Contrary to the usage of 1,024 as a cutoff point for several other epitope prediction models^8,11,15^, which is selected as the maximum length of sequences that the original ESM-2 model is trained on, the architecture supports much longer sequences. ESM-2 utilizes a rotary positional embedding (RoPE) encoding framework, and as RoPE allows for relative positional information, inference and modification of the model are not technically bound to set sequence lengths like traditional absolute positional embeddings^13^.

While some epitope prediction tools do not distinguish between linear and conformational epitopes, the difference in dataset size and prediction task (sequential vs sparse prediction) prompted us to distinguish between linear and conformational fine-tuning. We tuned linear and conformational versions of ESM-2 35M, 650M, and 15B. To illustrate the performance gains from including longer sequences, we also tuned several models on a version of the linear dataset which truncated sequences past 1,024 positions, and separately on a version which removed sequences longer than 1,024 altogether. Since several peer models did not fine tune, we also trained a model, 35m-trained, with identical architecture to ESM-2 35M but initialized with random weights. This 35m-trained model would serve as a baseline to highlight the performance of training on the architecture directly as opposed to fine-tuning ESM’s weights. A summary of all models discussed here is provided in (Supplementary Materials, Table 2).

In order to fine tune on the 15B version of ESM2, we incorporated PEFT-LoRA^17^ along with the Accelerate library^18^. The Accelerate library allowed us data parallelization across our GPUs, thus preventing OOM errors, especially in the case of the 48-layer 15B parameter model. Low-rank adaptation (LoRA)^19^ allows us to break up weight matrices of shape R^(M,N) into trainable low rank matrices of shape R^(M,low_rank) and R^(low_rank,N) and only update those low rank matrices. These low rank matrices are then multiplied together to restore their original shape, before being added back into the original matrices. Incorporating this PEFT-LoRA architecture serves to further reduce maximum memory consumption and allows the model to fit more aggressively to our epitope prediction task. Fine-tuning was performed on a server with 4×80GB A100 GPUs for 40 epochs beginning at a 1e-5 learning rate and 0.001 weight decay.

Batch size chosen as 1 to minimize peak memory consumption. PEFT-LoRA was initialized with ‘TOKEN.CLS’ task with r=8, lora_alpha=32, lora_dropout=0.1, and LoRA indicated for ‘query’, ‘key’, and ‘value’ matrices.

**Figure 1.**
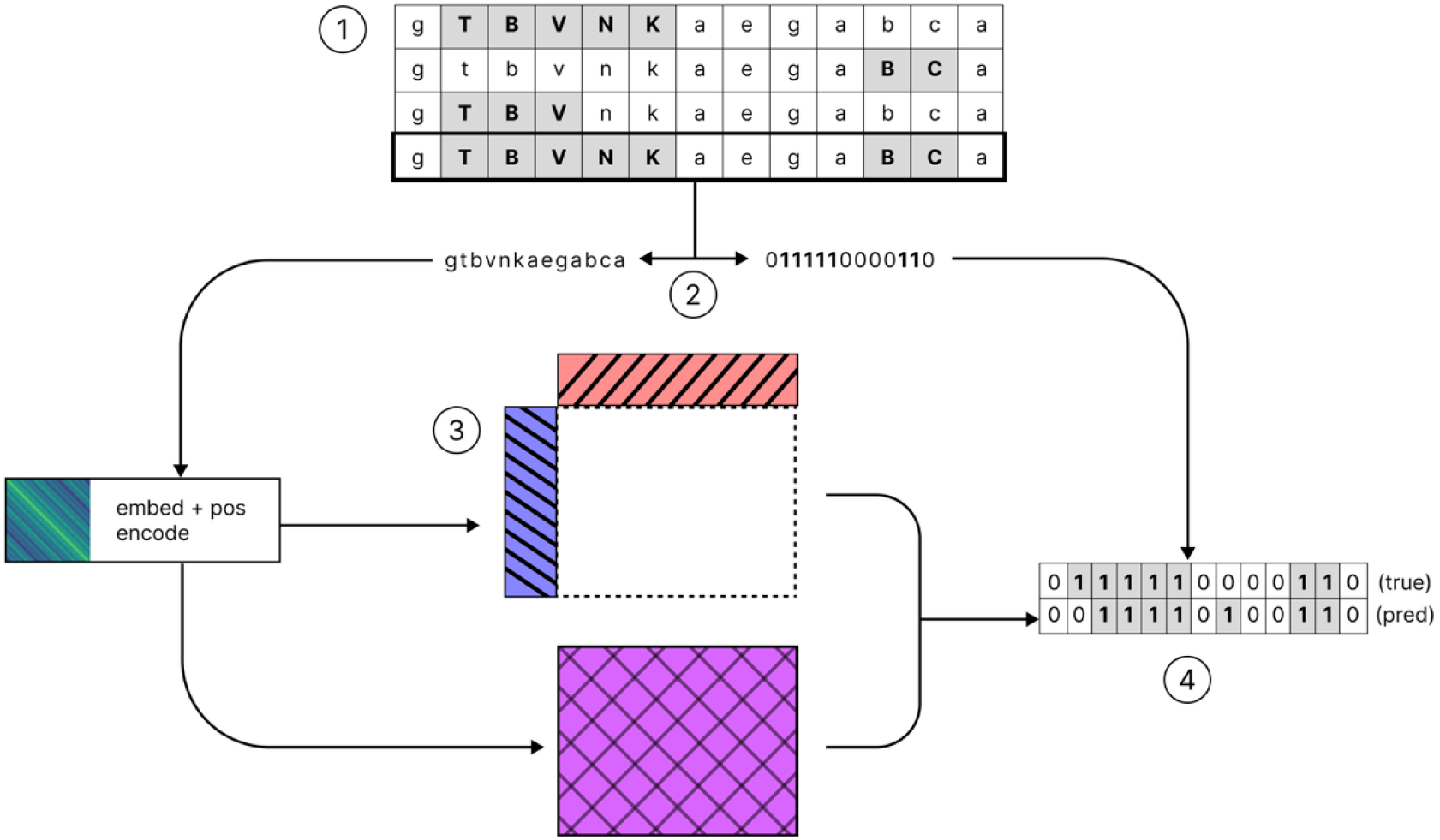
1) All epitopes of a given antigen are combined onto the antigen sequence. 2) The epitope annotation information is separated from the sequence; the un-annotated sequence is sent into embedding space and enters the model while the epitope positions are saved to calculate losses. 3) The model completes a forward pass across both our original weights and our low-rank matrices, whose outputs are summed together. 4) Softmax applied post-MLP used to calculate loss against annotated epitope positions generated previously. Only the low-rank matrices are directly updated in the backwards pass, and are then restored to their shape and summed with the original matrices.

### Comparison to Peer Models

To assess other state-of-the-art methods for epitope prediction, we tested our linear and conformational test set sequences on sequence-based and structure-based epitope prediction models. Models were assessed on the basis of global ROC-AUC, PR-AUC, and precision and recall at the thresholds provided by their respective publications.

BepiPred 3.0 and SEMA 1D were used to assess current sequence-based prediction, and DiscoTope 3.0 and ScanNet were used to assess current structure-based prediction.

Sequences were submitted to the online tools of BepiPred 3.0 and SEMA-1D. All models produced epitope scores per position, which were classified using their default thresholds for epitope classification: 0.1512 for BepiPred 3.0, and 0.361 for SEMA-1D.

To assess structural-based prediction models, we retrieved structures of our test set sequences from PDB. Of 566 linear test sequences, only 107 had crystal structures with all IEDB-identified epitope positions modeled, and AlphaFold structures were used for 380 of the remaining 441 antigens. 79 antigens had no suitable structure available, and were excluded from DiscoTope and ScanNet’s evaluation. DiscoTope’s online tool was used at its default threshold of 0.9 (identified as ‘moderate confidence’)^7^. ScanNet was assessed locally. The ScanNet repository did not provide a threshold for epitope classification, so a default threshold of 0.5 was used for epitope classification. Both models required alignment between crystal structures and IEDB antigens to be scored due to the inconsistency between their sequences.

Epitope classifications were compared across all approaches on linear and conformational sequences. EpiTune’s training produced checkpoints at every epoch, with both EpiTune 650M and 15B overfitting well before training completed. To avoid overfit checkpoints, the checkpoints used for model comparison were chosen based on each model’s best PR-AUC performance on the test set.

### Attention & Attribution Mapping

By utilizing only one model in its prediction pipeline, EpiTune also offers greater ease in interpretability, or analyzing the meaningfulness the model assigns to features. We demonstrate this by examining the long-distance (>1,024 residues) token-to-token relationships that ESM-2 not only supports, but reinforces when fine-tuned on the epitope prediction task. To better understand these relationships, we also fine-tuned with truncated and short sequences to reflect training techniques observed in the literature^8,9^. EpiTune-Linear-35m and ESM-2-35m were used as baselines to compare to EpiTune-Linear-35m-truncated, which kept all sequences but truncated past 1,024 residues, and EpiTune-Linear-35m-short, which only kept sequences <1,024 residues in length. These models were trained with the same starting parameters, including fixed RNG seeds, as the default EpiTune-Linear-35m to ensure consistency.

Once all models were trained, we compared model attention maps directly by plotting attention matrices for each model with PyTorch and Matplotlib. Since attention does not conclusively reveal the importance of information used to make predictions^20^, we also incorporated Captum’s LayerIntegratedGradients to explore the attribution of input importance, and LayerConductance to determine attribution on the attention layers. Using Layer Conductance^21^, we retrieved the attribution of the attention matrices per layer, and mapped them back onto our attention to get matrices of token-token interaction attribution. Normalizing across our heads dimension and summing attributions per token-token distance, we are able to visualize how local or distant token-token interactions contribute to the final prediction.

## Results

### Dataset

The linear and conformational datasets were comprised of 6,284 and 321 total antigens, respectively, after removing sequences longer than 1,724 residues. Figure 2 plots the distribution of antigen sequences by length. These datasets contained all positive human b-cell epitopes found on IEDB mapped onto antigens. However, mapping all epitopes onto their respective antigen sequences without negative-assay filtering results in approximately 5% of antigen sequences labeled as entirely epitope (Supplemental Fig 1A): this is primarily due to the large assay submission volume for certain sequences, such as SARS-CoV2 Spike glycoprotein^22^. Supplemental Figure 1B shows that our choice of removing epitopes with a greater amount of negative assays than positive ones served as an effective filter on overly saturated antigen sequences. Conformational datasets were not filtered.

**Figure 2.**
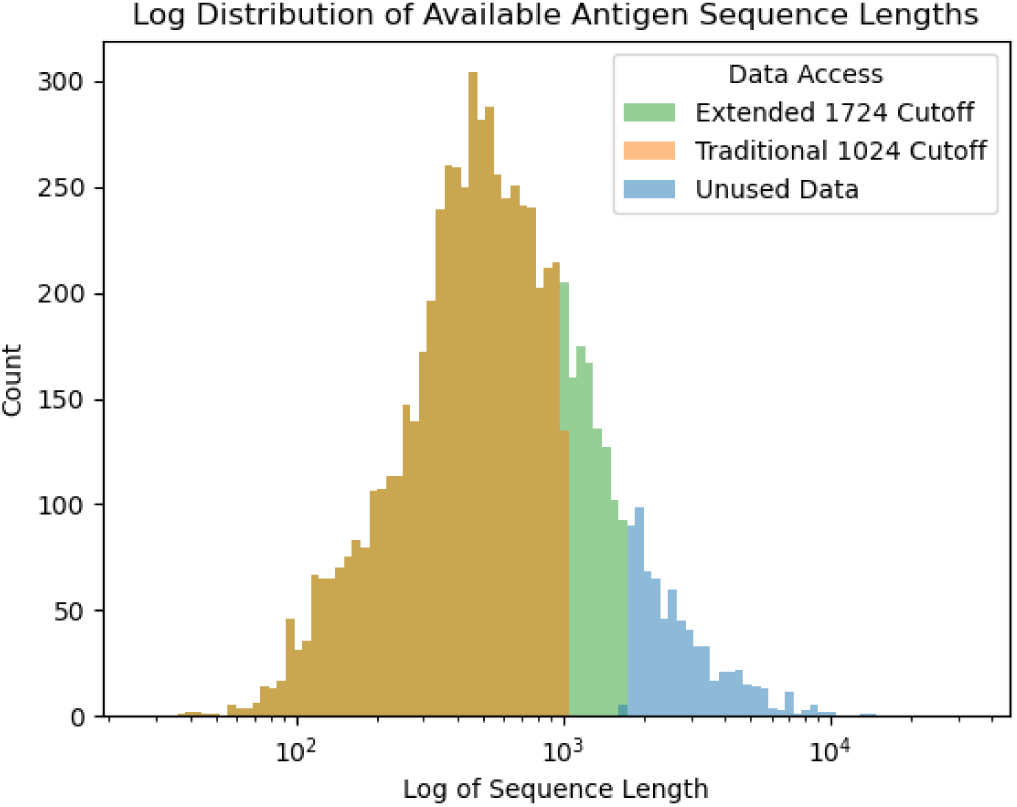
Distribution and segmentation of the available linear epitope sequences. Models are traditionally trained with sequence lengths below 1,024 residues. EpiTune was trained with sequence lengths up to 1,724 residues.

Separately, making use of ESM’s RoPE framework allowed us to add approximately ∼1,000 antigens to our linear training data (green section in Figure 2). This represented a significant increase in the portion of the training data available to our model, and allowed us to explore whether there are differences in model performance on sequences longer than ESM’s cutoff point. This added 39 sequences to the conformational dataset.

We created several modified datasets in order to tune different versions of EpiTune-Linear. In addition to our base EpiTune-Linear, we tuned short and truncated versions of EpiTune-Linear with their own datasets. The “short” versions of EpiTune only used sequences that were less than 1,024 residues in length (Figure 2, orange), and are meant to reflect the traditional 1,024 residue training cutoff. Consequently, the “short” test set contained only 477 sequences. The “truncated” versions were trained on all available sequences truncated past 1,024 positions, resulting in a truncated test set containing 629 sequences. The test set of standard EpiTune-Linear contains 566 sequences at a 1,724-length cutoff. Sequence limits are imposed before train/validation/test splitting, so test sets for the various EpiTune-Linear versions are not subsets of each other, and thus must be evaluated independently to prevent overlap in scoring. Finally, the conformational dataset yielded a final count of 32 test sequences.

### EpiTune Performance on Linear Epitopes

Table 1 lists the performance of the EpiTune-Linear models, peer models, and the 35m-trained model on linear epitopes from the test set consisting of 566 sequences. The 15B model showed significant improvement over 35M and 650M parameters. Precision increased 3% while recall increased 20.5%, indicating the 15B model is much better in positively identifying all epitope residues, and a smaller benefit in positively identifying residues as epitopes. Both contribute to an 18% improvement in PR AUC. By way of example, Figure 3 illustrates the Histone-binding protein RBBP4 epitope as reported in IEDB versus the EpiTune-Linear 35M and 15B predicted epitopes.

**Figure 3.**
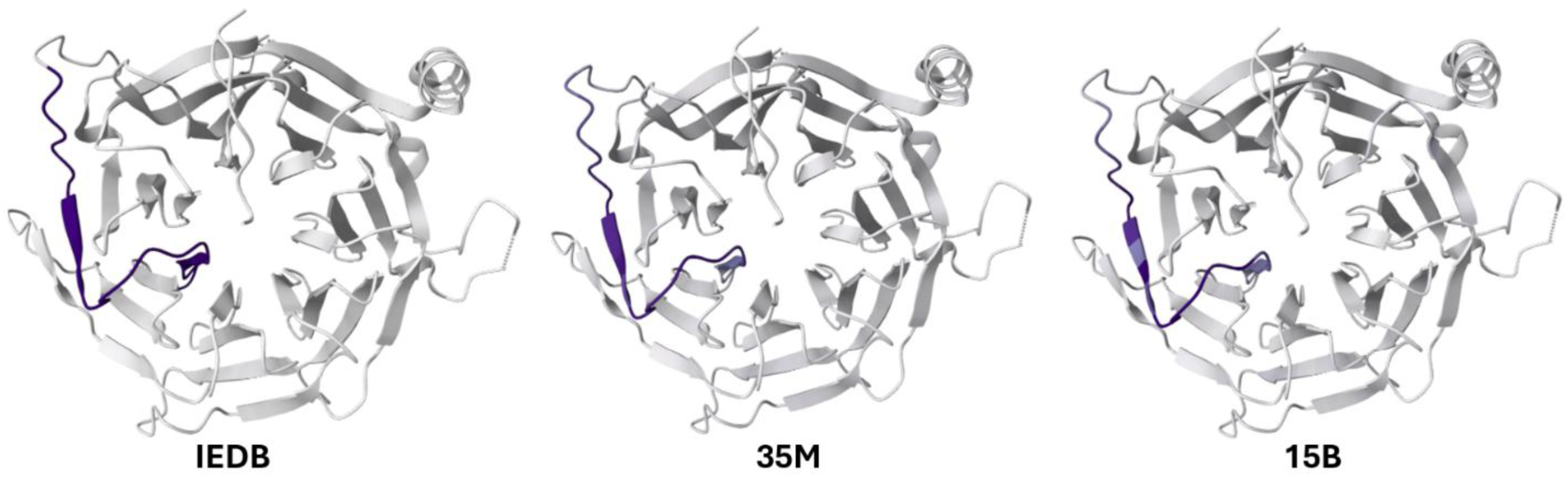
Histone-binding protein RBBP4 epitope (blue) identified experimentally and deposited in the IEDB (IEDB) compared to those predicted by the EpiTune-Linear 35M-parameter (35m) and 15B-parameter (15b) models.

**Table 1.** Performance of EpiTune and peer models on predicting linear and conformational epitopes. Each model was trained, tested, and validated with the appropriate data sets.

| Model | Precision | Recall | PR AUC | ROC AUC |
| --- | --- | --- | --- | --- |
| *Linear* |  |  |  |  |
| EpiTune-Linear 35M | 0.35 | 0.44 | 0.33 | 0.80 |
| EpiTune-Linear 650M | 0.27 | 0.43 | 0.25 | 0.75 |
| EpiTune-Linear 15B | 0.36 | 0.53 | 0.39 | 0.83 |
| 35m-trained | 0.08 | 0.99 | 0.08 | 0.50 |
| BepiPred 3.0 | 0.17 | 0.25 | 0.14 | 0.64 |
| SEMA 1D | 0.08 | 0.73 | 0.09 | 0.54 |
| DiscoTope 3.0 | 0.07 | 0.22 | 0.10 | 0.58 |
| ScanNet* | 0.06 | 0.96 | 0.06 | 0.50 |
| *Conformational* |  |  |  |  |
| EpiTune-Conf 35M | 0.06 | 0.22 | 0.05 | 0.62 |
| EpiTune-Conf 650M | 0.06 | 0.30 | 0.05 | 0.67 |
| EpiTune-Conf 15B | 0.12 | 0.22 | 0.08 | 0.74 |
| BepiPred 3.0 | 0.07 | 0.29 | 0.05 | 0.65 |
| SEMA 1D | 0.06 | 0.74 | 0.18 | 0.74 |
| DiscoTope 3.0 | 0.01 | 0.33 | 0.01 | 0.54 |
| ScanNet* | 0.00 | 0.00 | 0.00 | 0.40 |

\*ScanNet did not provide a cutoff, so we used 0.5 as our threshold.

EpiTune-Linear-15B performance compared to the published ROC AUC values of DiscoTope 3.0 and SEMA 1D. A head-to-head comparison using similar datasets significantly improves EpiTune performance. Overall, the models had low precision and moderate ROC AUC values when tested on our IEDB test set. We note that these values are overall lower than their reported values and might be the result of differences in training that do not align with our test set. However, when models were assessed on a test set of all positive epitope assays (Supplemental Table 1), rather than our filtered epitopes, each model’s performance was similar to published values.

The 35m-Linear-trained model highlights the inferior performance of training on a randomly initialized model with identical architecture to ESM-2-35m. This model’s ROC AUC and PR AUC metrics closely match those of the non-EpiTune models, strongly hinting at fine-tuning being necessary to achieve strong epitope prediction.

Table 2 compares the performance of each of EpiTune’s linear versions. The truncated versions offered the worst performance, with EpiTune-Linear-15B-truncated performing worse than the standard EpiTune-35m. Both the standard EpiTune-Linear versions achieved the best performance on their test set, with EpiTune-Linear-15b performing at nearly a 0.4 PR AUC. EpiTune-Linear-Short performed middlingly on both tests.

**Table 2.**
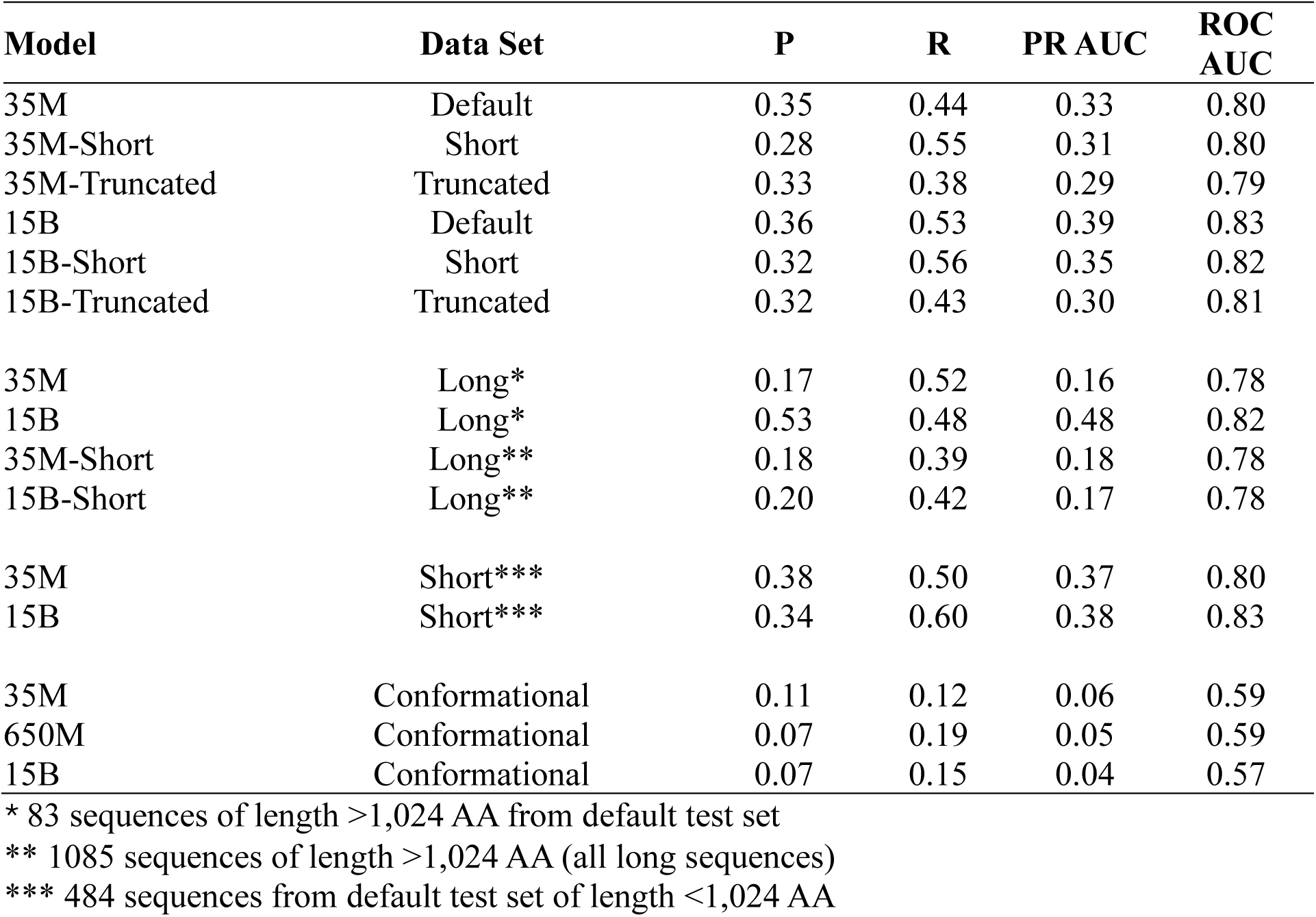
Alternative EpiTune-Linear Performances:

| Model | Data Set | P | R | PRAUC | ROC AUC |
| --- | --- | --- | --- | --- | --- |
| 35M | Default | 0.35 | 0.44 | 0.33 | 0.80 |
| 35M-Short | Short | 0.28 | 0.55 | 0.31 | 0.80 |
| 35M-Truncated | Truncated | 0.33 | 0.38 | 0.29 | 0.79 |
| 15B | Default | 0.36 | 0.53 | 0.39 | 0.83 |
| 15B-Short | Short | 0.32 | 0.56 | 0.35 | 0.82 |
| 15B-Truncated | Truncated | 0.32 | 0.43 | 0.30 | 0.81 |
| 35M | Long* | 0.17 | 0.52 | 0.16 | 0.78 |
| 15B | Long* | 0.53 | 0.48 | 0.48 | 0.82 |
| 35M-Short | Long** | 0.18 | 0.39 | 0.18 | 0.78 |
| 15B-Short | Long** | 0.20 | 0.42 | 0.17 | 0.78 |
| 35M | Short*** | 0.38 | 0.50 | 0.37 | 0.80 |
| 15B | Short*** | 0.34 | 0.60 | 0.38 | 0.83 |
| 35M | Conformational | 0.11 | 0.12 | 0.06 | 0.59 |
| 650M | Conformational | 0.07 | 0.19 | 0.05 | 0.59 |
| 15B | Conformational | 0.07 | 0.15 | 0.04 | 0.57 |

\* 83 sequences of length >1,024 AA from default test set
\*\* 1085 sequences of length >1,024 AA (all long sequences)
\*\*\* 484 sequences from default test set of length <1,024 AA

Table 2 also reports EpiTune-Linear performance on longer sequences, with base EpiTune-Linear assessing all long sequences in its validation set, and EpiTune-Linear-short assessing the long sequences excluded from its dataset (<1,724, >=1,024). Here, base EpiTune-Linear achieves decisively superior performance to its short-trained counterpart, with base EpiTune-Linear-15B scoring 0.48 PR-AUC, and EpiTune-Linear-15B-short achieving similar performance to both conformational and linear 35M counterparts (0.18 vs 0.17 and 0.16 PR-AUC). EpiTune-Linear-short’s moderate performance on short sequences and poor performance on long sequences appears to reflect a model stunted without the presence of long sequences in training, contrasting base EpiTune-Linear’s high performance for both tasks.

### EpiTune Performance on Conformational Epitopes

Table 1 lists the performance of the EpiTune-Conf models on conformational epitopes. All metrics were lower compared to their EpiTune-Linear counterparts. Precision and PR AUC were significantly reduced for the confirmational EpiTune predictions relative to linear.

The conformational dataset contains about 5% of the number of examples of the linear dataset. Furthermore, the conformational dataset has a lower average epitope ‘saturation’ per antigen, that is, the percentage of antigen sequence marked as an epitope (Supplemental Figure 1); increased class imbalance can lead to poorer performance of ML models^23^. A difference in dataset sizes is not sufficient to explain the difference in model performance, however. To mimic the small conformational dataset, several versions of EpiTune-Linear-35m were trained by selecting 350 sequences at random from the linear dataset. These models yielded PR AUC’s between 0.14-0.23, significantly higher than those produced by the EpiTune-Conf versions.

We also find that, despite much poorer prediction than their linear counterparts, increasing sizes of EpiTune-Conf models are able to learn the conformational epitope prediction task better. This, in turn, signals that larger models are necessary with this approach to correctly fit to the conformational epitope prediction space, and that fine-tuning our models brings them closer to this space. When assessing the performance of our linear models on the same conformational test set (Table 2), we see the opposite relationship: larger EpiTune-Linear models perform poorer. These observations remain true throughout each model’s training [Figure 4].

**Figure 4.**
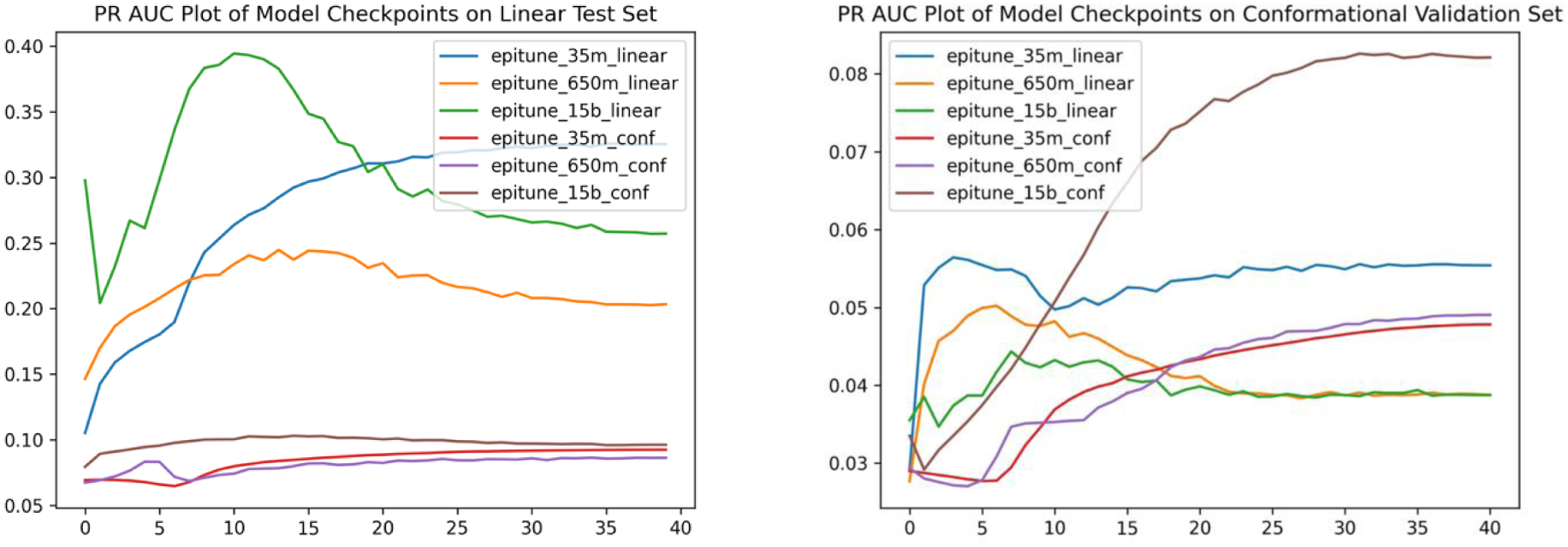
Performance of Linear and Conformational EpiTune Models on Linear and Conformational Test Sets. Note that, on their respective test sets, each model’s PR AUC scores increase as they are trained, but stay flat on the opposite test set.

Following this pattern, SEMA, which is trained specifically for conformational epitope prediction^9^, achieves the best performance on conformational prediction, and the worst performance on linear prediction (Table 1). This strongly suggests that prediction for linear and conformational epitopes are separate tasks, and should be considered separately.

### Attention & Attribution

EpiTune-Linear trained on non-truncated sequences has greater attention paid to distant points as opposed to the attention maps provided by the truncated EpiTune-Linear (Figure 5B). The difference between the two attention maps (in both attention gains and losses) follows the same sinusoidal-shaped pattern (Figure 5A) as the rotary positional embedding map. Despite the presence of rotary positional embedding, which helps generalize to longer sequences without explicit long-sequence training, actual long-sequence training impacts attention differently in both short and long-range token-token interactions.

**Figure 5.**
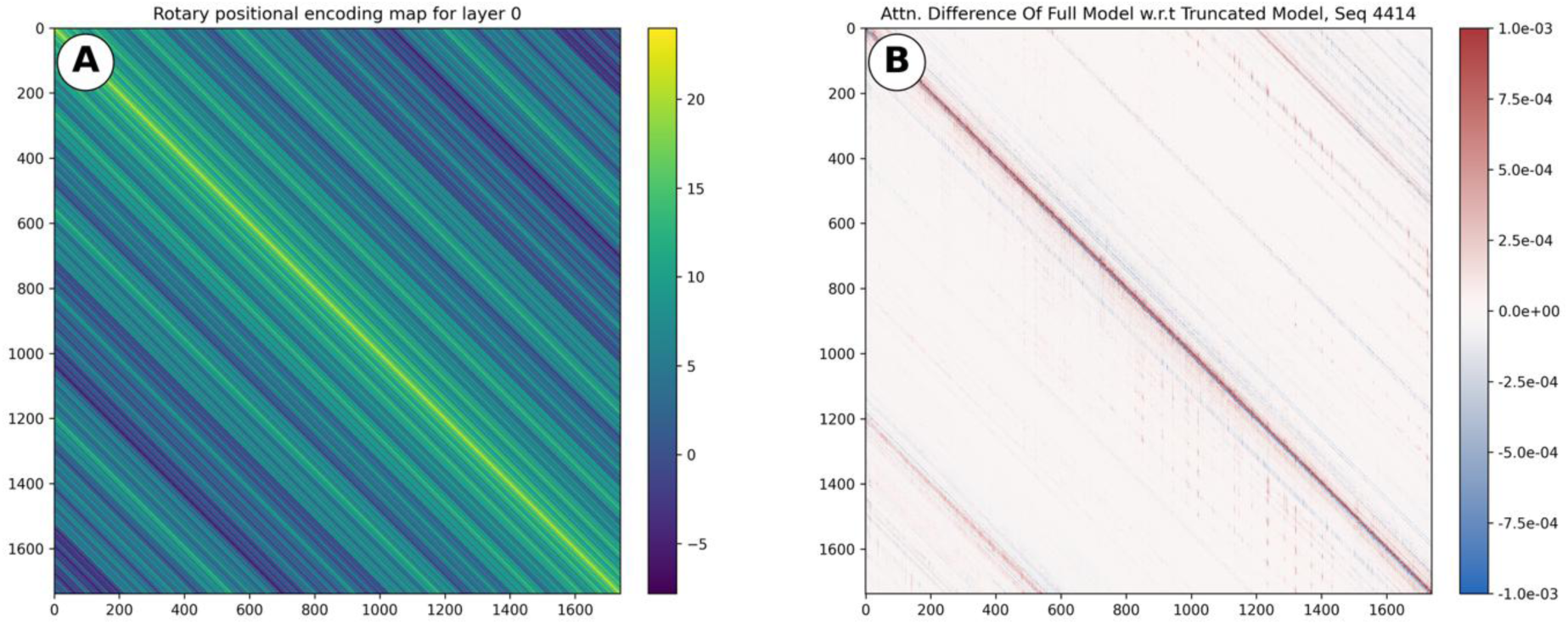
Subfigure A highlights the rotary positional embedding (RoPE) of a given position with respect to every other position, with brighter colors reflecting an increased embedding relevance score. Unsurprisingly, the rotary embedding scores are nonzero past the 1,024 truncation length that is often used as a cutoff for training epitope prediction models^8,9^. Figure 5B shows the difference between the attention scores of EpiTune-Linear-35m and EpiTune-Linear-35m-Truncated when predicting on a long (1,736AA) sequence. Attention scores were calculated across every head and layer. Positive values reflect where attention was increased in the standard EpiTune-35m-Linear model compared to the truncated model, and negative values reflect where attention was decreased relative to the truncated version. Note that the attention increases at distances further than the model was trained at following the rotary positional embedding pattern.

In all versions of EpiTune-35m-Linear we observed greater distant token-token attribution compared to the base ESM-2-35m. One explanation is that epitope prediction requires the synthesis of longer-range information than the protein feature prediction tasks used to train ESM-2. However, attributing directly to the Linear-35m model attention matrices produced high convergence deltas (∼0.5 average delta across all layers) for each EpiTune version, so the differences in attribution per layer for token-token interaction were assessed qualitatively and are not conclusive.

We explore how the models attribute to sequences of differing lengths in (Figure 6). These attributions had Layer Integrated Gradient convergence deltas close to 0. Here, we see clearer differences in model attribution due to differences in training sequence lengths. While all tuned models perform similarly on sequences <1,024 AA lengths, their attribution diverges markedly with the longer sequences. In all examples, standard EpiTune-Linear achieves superior performance, despite sequence overlap with the training sets of EpiTune-Linear-Short and EpiTune-Linear-Truncated. We also see that the standard EpiTune-Linear is more conservative with the residues that it attributes to in longer sequences, whereas both EpiTune-Linear-Short and EpiTune-Linear-Truncated are less certain. In all cases, the full EpiTune-Linear-35m has the best performance in predicting on the selected sequences.

**Figure 6.**
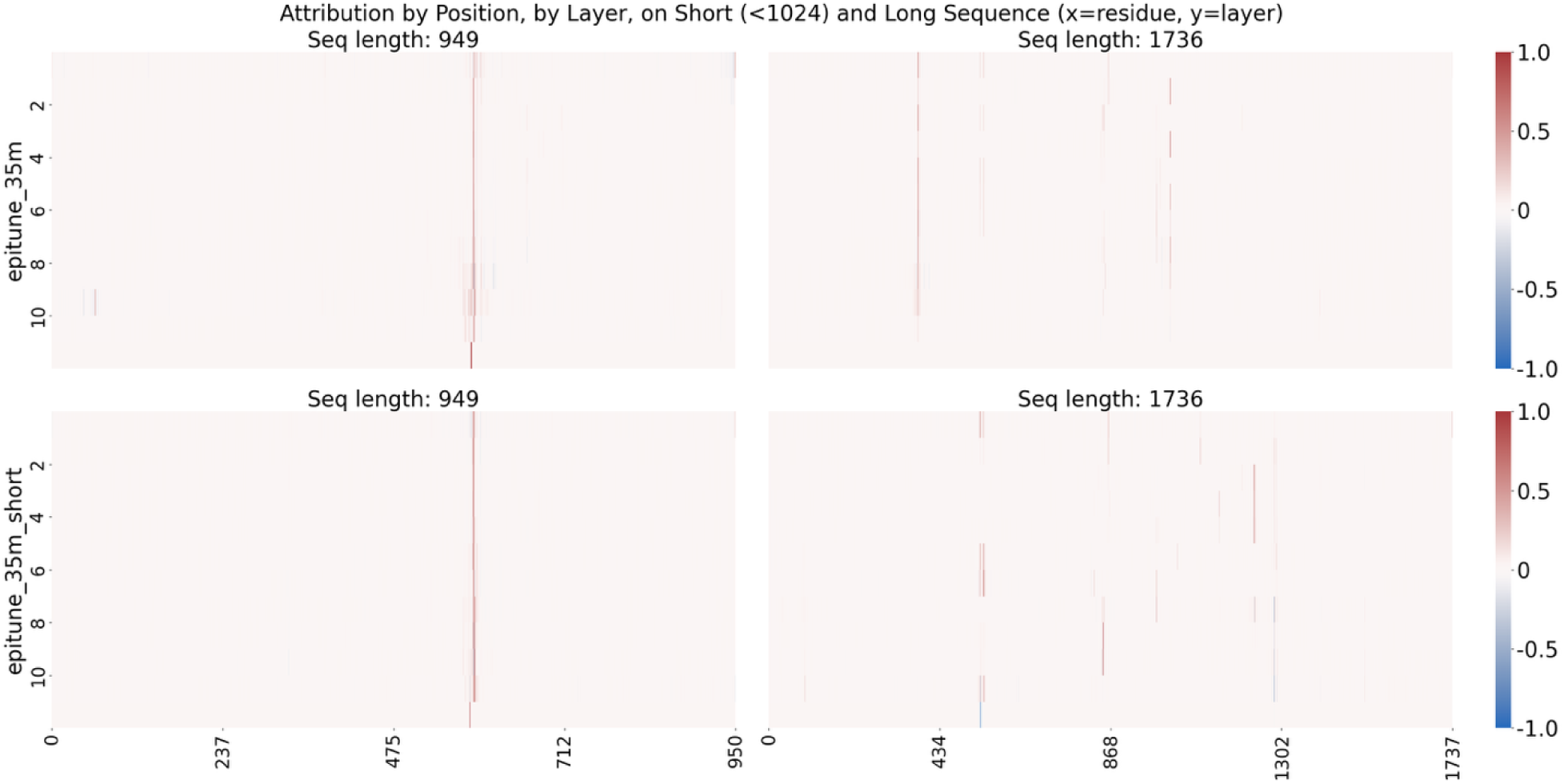
Comparison of attributions of EpiTune-35m-Linear variants on short and long sequences. Note the consistency of attribution in the short sequence, and difference of attribution in the longer sequence. For a full comparison of attribution across all EpiTune-35m-Linear variants across more sequences, refer to Supplementary Materials Figure 2.

## Discussion

EpiTune-Linear demonstrates the capability of a pure transformer model to accurately predict linear epitopes through the direct tuning of ESM-2. In contrast, several state-of-the-art models incorporate ESM-2 into their model architecture primarily as an embedding device alongside a biologically interpretable model. This tempting approach has clear benefits:

-**Directly training transformer models is computationally expensive,** as they are less time and space efficient than other models^24^. Inference with transformer models is much cheaper, so some of their benefits are still leveraged by using them as embedding tools.

-Researchers often incorporate concepts in models known to physically reflect the biology of the epitope or protein, such as SEMA’s n-glycosylation model^9^. Using multiple models enables researchers to easily **incorporate pieces of information besides the sequence’s embeddings**, such as SASA/Shapley values^7^. These additional values are expected to improve prediction as they reflect protein rules in the real world.

**-Starting with a new (randomly initialized) model allows the model to fit strictly to the epitope prediction task.**

EpiTune’s approach allows us to solve many of the computational challenges of training, and highlights that the assumptions around feature importance are better determined by a model trained on the feature space.

We were able to overcome the hurdle of computationally expensive training with PEFT-LoRA^17^ and make the models and training materials available on our GitHub repository^25^. Transformers are superior to earlier models for sequential prediction tasks in high-dimensional latent spaces that involve predictions based on long-distance^24^. Previous architectures struggle with sequential long-distance prediction due to vanishing or exploding gradients^26^.

Other models make use of information besides the antigen sequence in predicting epitopes. Our work suggests that, at least for linear epitope prediction, these features are encoded in, and can successfully be learned from, the antigen sequence alone. Framing epitope prediction in biologically interpretable terms is unnecessary for performant prediction^27^, and assumes that human interpretable/physically reflective intermediates, such as antigen structure, are important in the epitope prediction task. Transformer models move information through circuits^28^, or sets of neurons responsible for completing the whole or part of a specific task. These pathways are made to minimize loss, ultimately increasing predictive power, and are not required to be decomposable into human interpretable concepts^28^. Additionally, because transformers have been shown to develop reasoning capabilities within their architecture while being trained on a task, a single model can facilitate the best performance. We also believe interpretability is possible in these models^28^.

Transfer learning is an effective tool^29^ and should be leveraged rather than having the model learn protein rules alongside epitope rules. In fact, 35m-Linear-trained [Table 1] essentially proves this point: with identical training and test data, architecture, training strategy and initial hyperparameters to EpiTune-Linear-35m, the randomly initialized 35m-Linear-trained performs much worse. This is not surprising, as epitopes are proteins, and benefit from the model first conforming to protein space and being moved towards the epitope direction, rather than having to move simultaneously towards the protein and epitope directions.

Although we demonstrated matching performance on the unfiltered test set (Supplementary Table 1), we were largely unable to recreate other publications’ claimed ROC AUC with our filtered test set (Table 1). While we cannot expect the same ROC AUC, we would expect similar or even slightly higher ROC AUC as our test sequences were chosen at random, thus allowing for overlap if the test sequences were present in another model’s training set.

One possible explanation for this is the difference between our training set formats. EpiTune utilizes the entire antigen sequence from IEDB, whereas the structure-based approaches (DiscoTope and ScanNet) only use the portion of the antigen for which a structure exists. Available PDB structures sometimes differ in residues reported from its IEDB antigen sequence due to the presence of isoforms and unsolved regions. Finally, of the test set, only 107 out of 566 antigens structures had crystal structures available in PDB^30^ that contained every epitope position on the antigen all other predictions used PDB structure files downloaded from AlphaFoldDB^31–33^.

The compared sequence-based models, such as SEMA-1D and BepiPred 3.0, follow the training cutoff point of ESM-2 near 1,024 residues. Both models follow a truncation-based approach, with SEMA truncating at 1,022 positions^9^, and BepiPred 3.0 at 1,023^8^. This cutoff point is not explicitly mentioned in SEMA’s publication, but is found within their GitHub repository. We observed that truncation-based cutoff approaches yielded the worst performance across the alternative versions of EpiTune-Linear (Table 2).

Another explanation of the discrepancy of performance across models is how the over-annotation of epitopes on certain antigen sequences bleed into model training sets. Interestingly, ROC_AUC and PR_AUC performance was roughly aligned with each model’s published values when used on the unfiltered (all positive assay epitopes present, no removal of epitopes if negative results were the assay majority) dataset with the exception of SEMA-1D (Supplemental Table 1). The EpiTune-Linear dataset filtered out epitopes where IEDB indicated that the epitope region returned a greater amount of negative assay results than positive assay results (Supplemental Figure 1). DiscoTope did not filter out by IEDB assay agreement, but seemingly also dealt with epitope ‘over-saturation’ on an antigen, filtering out antigens with >75 epitope positions annotated^7^.

### Linear and Conformational Prediction

While EpiTune-Linear surpassed other linear epitope prediction models, both sequence and structure based, EpiTune-Conf struggled in conformational epitope prediction. Our results highlighted that this discrepancy not only arises from a dearth of conformational data, but from the fact that conformational prediction appears to be a separate task from linear prediction altogether. Though not included here, early designs of EpiTune incorporating transfer learning between linear and conformational models performed worse than training on the tasks separately. This observation is consistent with the findings in CALIBER^34^. Because several models (e.g. DiscoTope, BepiPred) still make no distinction between performance on linear or conformational epitopes, future work should assess both types of epitopes, and consider training for these tasks separately.

If conformational prediction is a more difficult task than linear prediction, then the fairly low rank PEFT-LoRA settings EpiTune-Linear used may result in poorer performance in tuning EpiTune-Conf. Further work should be done to determine the optimal hyperparameters for low rank adaptation in conformational prediction, and to what extent hyperparameter choice and low rank adaptation impact the performance of linear versus conformational epitope prediction tasks on differing model sizes.

### Applicability of RoPE in Long Sequence Prediction

We observe that there are differences in how the versions of EpiTune perform (Table 2) and think (Figure 6, Supplementary Materials Figure 2) with longer sequence lengths. We see that the learned epitope prediction on shorter sequences, as represented by the short and truncated versions, does not translate to longer sequences. The ability to handle longer sequences comes with a moderate performance tradeoff on shorter sequences, which may have several implications for fine-tuning models on the epitope prediction task.

The ESM-2 publication^12^ states that RoPE was added to allow the model to extrapolate beyond the context window it was trained on, but does not provide measurements of performance on longer sequences. ESM incorporates a mixed strategy with rotary embeddings, using absolute embeddings for the first 1,024 tokens before switching to RoPE. In the case that the RoPE architecture employed in ESM-2 allows the model to perfectly generalize to longer sequences as if trained with absolute embeddings, a difference in performance between long and short sequences implies that the epitope prediction task is fundamentally different depending on sequence length. This would explain both the failure to generalize from the short models, as well as the higher performance of short models on short sequences, and the lower performance of the standard models on shorter sequences (Table 2).

In the more likely scenario that RoPE does not perform perfectly, this discrepancy is explained by the same reason our fine-tuning was superior to training a model from randomly initialized parameters: RoPE could not successfully generalize to the longer sequence space, and so, during fine-tuning, our model has to learn both the long protein space and the epitope space. This calls into question the translatability of applying long-sequence RoPE with minimal performance degradation^13^ to the biology domain. Alternatively, the RoPE settings used by ESM-2 may be inappropriate for the epitope prediction task. Further work must be done to determine whether overall protein prediction, or epitope prediction specifically, fails to generalize from the short-trained models to longer sequences.

