## Supplemental Material for "EpiTune: An Accurate Epitope Prediction Model with Mechanistic Insights"

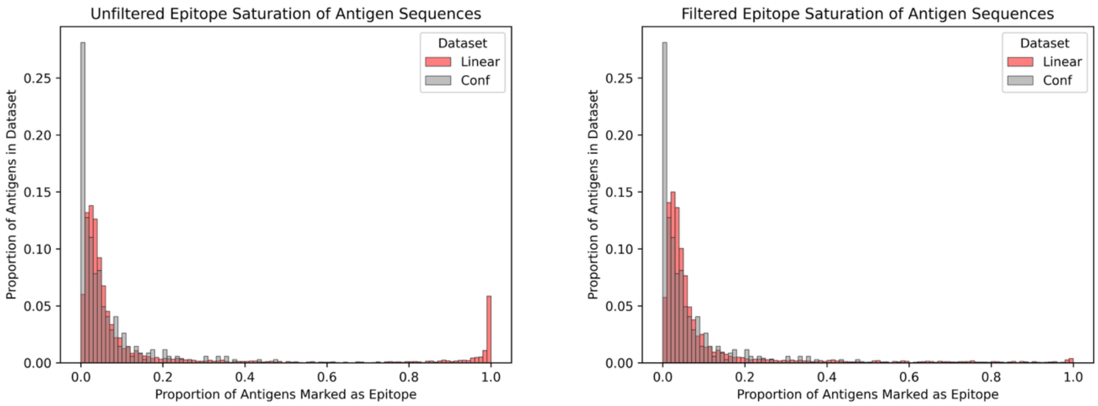


**SM Figure 1**. Distribution of ‘epitope saturation’, or the proportion of an antigen that is marked as an epitope. The left plot (Item 1 A) shows the distribution of linear epitopes and antigens before filtering out epitopes with majority negative assay results. The right plot highlights the dataset distribution after filtering out these epitopes.

**SM Table 1**.Table comparing performance of EpiTune-Linear-Positive versus other state-of-the-art tools on linear epitopes with unfiltered dataset. The dataset used to create the test and training sets, as well as to tune EpiTune-Linear-Positive, consisted of **all** positive b-cell human-hosted epitopes within IEDB. Note that the ROC_AUC and PR_AUC metrics are much higher here than in the Table 1, and are close to each model's published metrics.

| **Model** | **Precision** | **Recall** | **PR AUC** | **ROC AUC** | **Claimed** |
|---|---|---|---|---|---|
| BepiPred 3.0 | 0.287 | 0.300 | 0.265 | 0.682 | 0.663-0.771 |
| SEMA 1D | 0.149 | 0.590 | 0.167 | 0.578 | ~0.75 |
| DiscoTope 3.0 | 0.065 | 0.182 | 0.187 | 0.734 | ~0.8 |
| ScanNet* | 0.056 | 0.001 | 0.083 | 0.544 | -- |
| EpiTune 35M Positive | 0.449 | 0.461 | 0.442 | 0.738 | -- |
| EpiTune 650M Positive | 0.633 | 0.393 | 0.489 | 0.743 | -- |
| EpiTune 15B Positive | 0.459 | 0.459 | 0.537 | 0.788 | -- |

*ScanNet did not provide a cutoff so we used >0.5 as our threshold.

SM Table 2. (Can be moved into main section) Model Summary

| Model | Architecture | Medium | Sequence Length Limit | Truncation? | Fine-Tuned? |
|---|---|---|---|---|---|
| BepiPred 3.0 | ESM-2 + FFNN | Sequence | 1,023 | Yes | No |
| SEMA 1D | ESM-2 + SaProt | Sequence | 1,022 | Yes | Yes |
| DiscoTope 3.0 | ESM-2 + CatBoost | Structure | None | No | No |
| ScanNet | ScanNet | Structure | None | No | No |
| EpiTune Linear | ESM-2 | Sequence | 1,724 | No | Yes |
| EpiTune Linear Short | ESM-2 | Sequence | 1,024 | No | Yes |
| EpiTune Linear Truncated | ESM-2 | Sequence | 1,024 | Yes | Yes |
| 35m-Linear | ESM-2 | Sequence | 1724 | No | No |
| EpiTune Conf | ESM-2 | Sequence | 1724 | No | Yes |

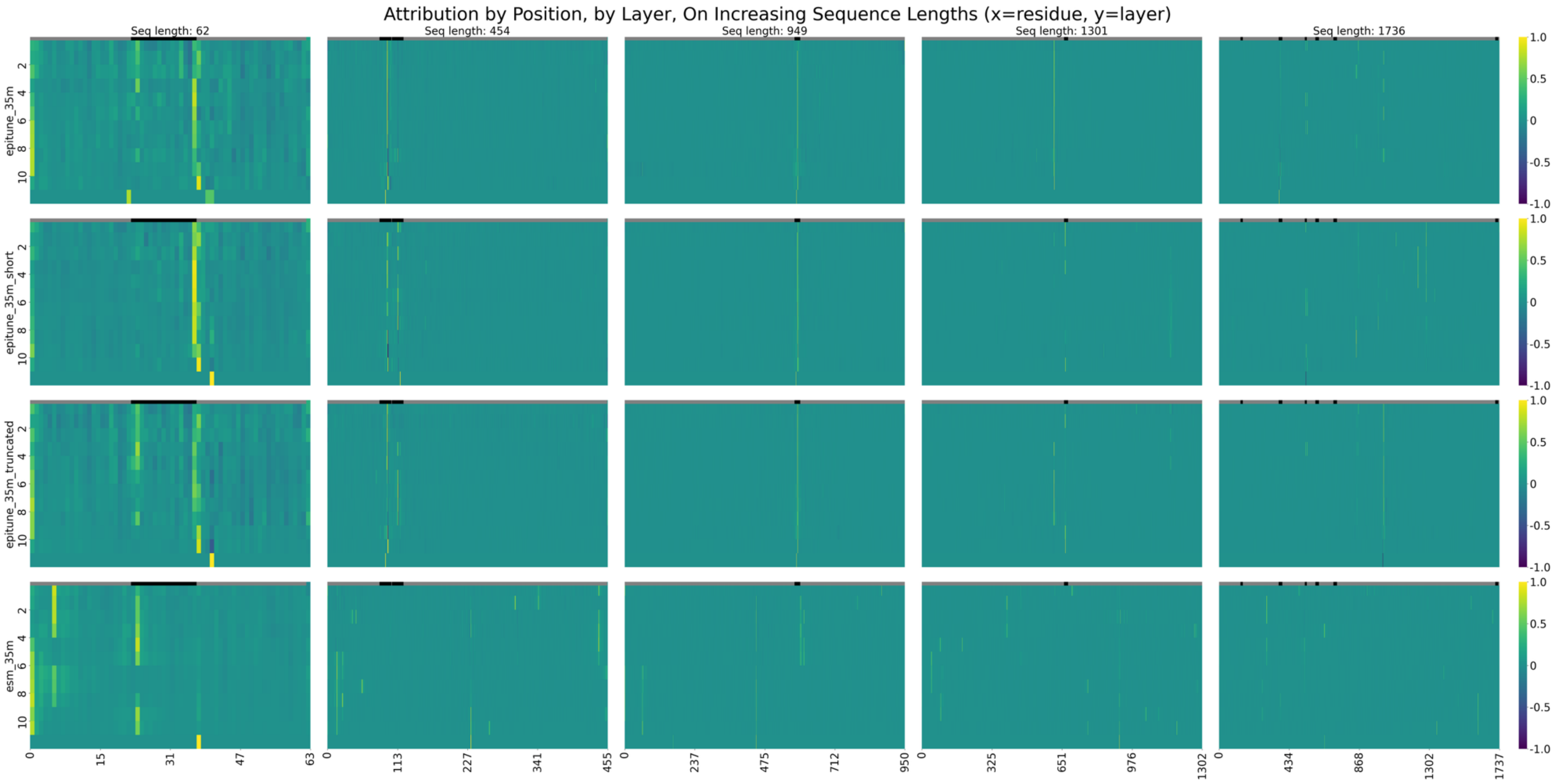


Supplementary Materials, Figure 2. Top row is standard EpiTune-Linear-35m, second is EpiTune-Linear-35m-short, third is EpiTune-Linear-35m-truncated, fourth is ESM2-35m. The columns are attributions on sequences of length 60/450/950/1300/1750. Note the consistency of attributions in each antigen sequence until the >1,024 sequence length point.
